# Hybrid incompatibility caused by an epiallele

**DOI:** 10.1101/099317

**Authors:** Todd Blevins, Jing Wang, David Pflieger, Frédéric Pontvianne, Craig S. Pikaard

**Author notes:** Present address: Université de Strasbourg, CNRS, IBMP UPR 2357, F-67000 Strasbourg, France. Present address: CNRS UMR5096, Laboratoire Génomes et Développement des Plantes, Université de Perpignan Via Domitia, Perpignan, France. Co-corresponding authors: Craig S. Pikaard, Todd Blevins. These authors contributed equally (Todd Blevins, Jing Wang).

## Abstract

Hybrid incompatibility resulting from deleterious gene combinations is thought to be an important step towards reproductive isolation and speciation. Here we demonstrate involvement of a silent epiallele in hybrid incompatibility. In *Arabidopsis thaliana* strain Col-0, one of the two copies of a duplicated histidine biosynthesis gene, *HISN6B* is not expressed, for reasons that have been unclear, making its paralog, *HISN6A* essential. By contrast, in strain Cvi-0, *HISN6B* is essential because *HISN6A* is mutated. As a result of these differences, Cvi-0 × Col-0 hybrid progeny that are homozygous for both Col-0 *HISN6B* and Cvi-0 *HISN6A* do not survive. We show that *HISN6B* is not a defective pseudogene in the Col-0 strain, but a stably silenced epiallele. Mutating *HISTONE DEACETYLASE 6 (HDA6)* or the cytosine methyltransferase genes, *MET1* or *CMT3* erases *HISN6B’s* silent locus identity in Col-0, reanimating the gene such that *hisn6a* lethality and hybrid incompatibility are circumvented. These results show that *HISN6*-dependent hybrid lethality is a revertible epigenetic phenomenon and provide additional evidence that epigenetic variation has the potential to limit gene flow between diverging populations of a species.

**Significance statement:** Deleterious mutations in different copies of a duplicated gene pair have the potential to cause hybrid incompatibility between diverging subpopulations, contributing to reproductive isolation and speciation. This study demonstrates a case of epigenetic gene silencing, rather than pseudogene creation by mutation, contributing to a lethal gene combination upon hybridization of two strains of *Arabidopsis thaliana*. The findings provide direct evidence that naturally occurring epigenetic variation can contribute to incompatible hybrid genotypes, reducing gene flow between strains of the same species.

## Introduction

Mutations that accumulate in separate subpopulations of a species can facilitate reproductive isolation by engendering hybrid incompatibility, a reduction in fitness observed among hybrid progeny at the F1 or F2 generations (1-3). Bateson, Dobzhansky and Muller (4-6) independently proposed that gene mutations made benign by compensatory mutations in interacting genes within isolated subpopulations prove deleterious when subpopulations or strains interbreed, thereby contributing to speciation by reducing gene flow. Examples include the lethal consequences of a *Saccharomyces cerevisiae* protein directing improper splicing of essential *S. bayanus* mRNAs in *S. cerevisiae* × *S. bayanus* hybrids (3) or the hybrid necrosis that results from expression of incompatible innate immune receptors in *Arabidopsis thaliana* (1). Lynch and Force envisioned an alternative scenario by which hybrid incompatibilities might arise, as a result of gene duplication (7). Gene duplication initially results in gene redundancy, thereby relaxing constraints on sequence and functional divergence, but mutations most often transform one paralog into a nonfunctional pseudogene (8-10). Lynch and Force recognized that asymmetric resolution of gene duplicates in this manner could result in either paralog becoming the sole operational copy in different subpopulations of a species, such that hybrid progeny inheriting non-functional alleles of both subpopulations would suffer reduced fitness (7).

Non-mutational (epigenetic) gene silencing could potentially contribute to hybrid incompatibility via the scenario envisioned by Lynch and Force. In plants, silent epialleles segregating in Mendelian fashion can be stably inherited over many generations, and are known to affect a number of well-studied traits, including flower morphology (11, 12) and fruit ripening (13). Silent epialleles are inherited via the perpetuation of repressive chromatin states, and a number of plant chromatin modifying enzymes are known to participate in epigenetic inheritance, including the Rpd3-like histone deacetylase, HDA6, the ATP-dependent chromatin remodeler, DDM1, and the DNA methyltransferases, MET1 and CMT3 (14-24). These enzymes are important for maintaining patterns of cytosine methylation following each round of DNA replication, such that newly replicated daughter strands of DNA inherit the methylation status of mother strands.

Multicopy transgenes frequently become methylated and silenced, particularly when inserted into the genome as tandem, inverted repeats that can give rise to doublestranded RNAs (25). Such double-stranded RNAs can be diced into short interfering RNAs (siRNAs) that guide the cytosine methylation of homologous DNA sequences, a process known as RNA-directed DNA methylation (RdDM) (26, 27). Known cases of silencing that involve duplicated endogenous genes, such as the phosphoribosylanthranilate isomerase (*PAI*) or folate transporter (*AtFOLT*) genes of *Arabidopsis thaliana*, resemble transgene silencing in that complex sequence rearrangements coincide with duplication of a new gene to create a novel (non-ancestral) locus engendering siRNA production and homology-dependent DNA methylation (28, 29). Silencing of the ancestral *AtFOLT1* gene, directed by siRNAs derived from the novel *AtFOLT2* locus present in some strains, causes reduced fertility in strain-specific hybrid combinations (28). This interesting case study has shown that naturally occurring RdDM, involving a new paralog that inactivates the ancestral paralog *in trans*, can be a cause of hybrid incompatibility.

Here, we demonstrate an epigenetic basis for a previously identified case of hybrid incompatibility (30) involving a gene pair, *HISN6A and HISN6B*. The mechanistic basis for *HISN6* hybrid incompatibility was previously unknown. We show that the *HISN6B* gene of strain Col-0, and hundreds of other *A. thaliana* strains, is hypermethylated in its promoter region and is epigenetically silent, making its paralog, *HISN6A* essential. Inheritance of the silent *HISN6B* epiallele requires HDA6, MET1, and CMT3-dependent cytosine methylation, but is unaffected by mutations disrupting RdDM. Although methylated *HISN6B* epialleles can be stably inherited for at least 30 generations, they can revert to an active state if the epigenetically silent state is erased by passage through an *hda6* mutant background. This allows *HISN6B* to now rescue *hisn6a* null mutations in strain Col-0 and to restore compatibility with strain Cvi-0, in which *HISN6A* is defective due to an internal deletion. Collectively, our results demonstrate that HISN6-dependent hybrid lethality is a previously unrecognized epigenetic phenomenon.

## Results

*HISN6A* and *HISN6B* are duplicated, paralogous genes (Figures 1A, S1) encoding histidinol-phosphate aminotransferase, which converts imidazole-acetol phosphate to histidinol-phosphate in the histidine biosynthesis pathway (31, 32) (Figure 1B). The gene duplication occurred after *A. thaliana* diverged from a common ancestor with *A. lyrata* (33, 34), resulting in a segment of Chromosome 1 containing *HISN6B* (At1g71920) becoming duplicated on Chromosome 5, yielding the *HISN6A* locus, At5g10330 (32, 35). Although *HISN6A* and *HISN6B* coding regions are 100% identical at the amino acid level, homozygous *hisn6a* null mutations are embryo lethal in strain Col-0 because *HISN6B* is not expressed (Figure 1B, C)(30, 31). This differential expression of *HISN6A* and *HISN6B* can be observed by RT-PCR amplification followed by digestion with *Rsa* I, which cuts only *HISN6A* cDNA (Figure 1C). In wild-type (WT) Col-0, only *HISN6A* is expressed (Figure 1D). However, in *hda6-6* or *hda6-7 mutants* (36, 37), *HISN6B* is also expressed (Figure 1D), which is also true in mutants for *MET1* or *CMT3*, which encode enzymes responsible for cytosine maintenance methylation in the CG and CHG sequence contexts (38), respectively (Figures 1D, S2A). By contrast, *HISN6B* is not derepressed in mutants deficient for RNA-directed *de novo* cytosine methylation, such as *nrpd1-3* (Pol IV), *nrpe1-11* (Pol V) or *drm2* (Figure 1D, S2A).

**Figure 1.**
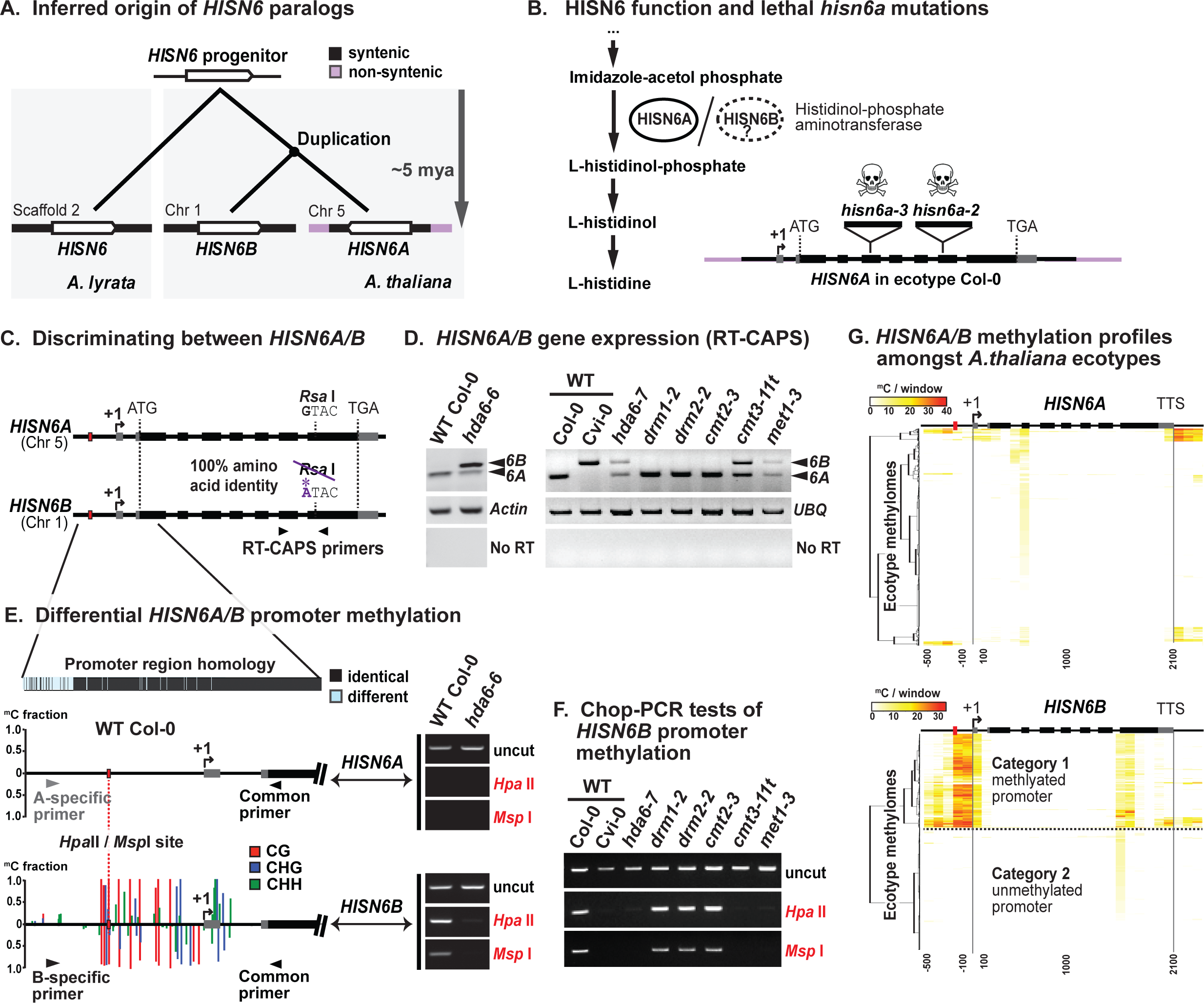
HDA6, CMT3 and MET1 silence *HISN6B* via promoter region DNA methylation. **(A)** Inferred phylogenetic origin of *HISN6* paralogs based on synteny between *HISN6B* flanking regions on *A. thaliana* chromosome 1 and sequences including the single copy *HISN6* gene of *Arabidopsis lyrata* scaffold 2 (see Figures S1A and B). **(B)** Left: HISN6A/B protein function in histidine biosynthesis. Steps upstream of Imidazole-acetol phosphate are omitted. Right: Gene structure of *HISN6A* in strain Col-0: untranslated regions (UTRs), exons and T-DNA insertion positions in mutant alleles are indicated by grey boxes, black boxes and inverted triangles, respectively. **(C)** Reverse transcription - cleaved amplified polymorphic sequence (RT-CAPS) assay for discrimination of *HISN6A* and *HISN6B* mRNAs using primers flanking a polymorphic RsaI site present only in *HISN6A* **(D)** *HISN6A/B* expression analysis via RT-CAPS in *hda6-6*, *hda6-7*, *drm1-2*, *drm2-2*, *cmt2-3*, *cmt3-11t*, *met1-3* mutants compared to wildtype (WT) Col-0 or Cvi-0. *Actin* and ubiquitin (*UBQ*) reactions serve as loading controls. Reactions omitting reverse transcriptase (no RT) are controls for genomic DNA contamination. **(E)** Analysis of DNA methylation in *HISN6A and HISN6B* promoter regions. Bar plots show WT Col-0 methylation profiles, color-coded by sequence context (CG, CHG and CHH), tabulated as fractional cytosine methylation (y-axis), based on methylome data of Stroud et al. (39). No methylation was detected in the *HISN6A* promoter. Gel images show Chop-PCR assays of cytosine methylation status at a *HISN6A/B* promoter *Hpa*II/*Msp*l site (red dotted line) in wild-type Col-0 or the *hda6-6* mutant. Reactions omitting restriction enzymes (uncut) demonstrate equivalent DNA input. *HISN6A/B* primer specificity was verified by sequencing PCR products **(F)** Analysis of *HISN6B* promoter methylation in the mutant series of panel D, using Chop-PCR **(G)** Hierarchical clustering of cytosine methylation profiles for *HISN6A* (top) or *HISN6B* (bottom) genes of 892 natural accessions (strains) of *A. thaliana*. Methylated cytosines, based on data from Kawakatsu et al. (40) were tallied within 100-bp, nonoverlapping windows starting 500 bp upstream of the transcription start site (+1) and stopping 300 bp downstream of the transcription termination site (TTS).

*HISN6* promoter CG and CHG methylation is easily assayed using Chop-PCR, a test in which genomic DNA is first digested (chopped) with methylation-sensitive restriction endonuclease *Hpa*II or *MspI* prior to PCR amplification of a region that includes a *Hpa*II/*MspI* recognition site, CCGG. In our tests, we assayed a *Hpa*II/*Msp*I site in the promoter region (Figure 1E, red dotted line), where Col-0 methylome data (39) show that dense CG and CHG methylation and scattered CHH methylation occurs in *HISN6B* but not in *HISN6A* (see Figure 1E gene diagrams). Using the Chop-PCR assay, PCR products were detected for *HISN6B* (Figure 1E, bottom right) but not for *HISN6A* (Figure 1E, top right), indicating CG and CHG methylation of the *HISN6B* promoter CCGG site, but not of the corresponding *HISN6A* site. *HISN6B* promoter CG and CHG methylation was lost in *hda6-6*, *met1-3* or *cmt3-11t* mutants, but not in Pol IV (*nrpd1-3*) or Pol V mutants (*nrpe1-11*) or mutants defective for the DNA methyltransferases DRM1, DRM2 or CMT2 (Figures 1E, F, S2B, S2C). Although 24-nt siRNAs matching the *HISN6B* promoter were detected by RNA-seq (Figure S2D)(14), their absence in the *nrpd1-3* mutant was not correlated with *HISN6B* reactivation. Collectively, the data of Figures 1D-F reveal that *HISN6B* silencing in Col-0 correlates with HDA6, MET1 and CMT3-dependent CG and CHG methylation. In strain Cvi-0, in which *HISN6A* has suffered a deletion mutation, *HISN6B* lacks promoter methylation (Figure 1F), and is expressed (Figure 1D). Analysis of publically available methylome data (40) reveals *HISN6B* promoter hypermethylation in 43% (387 of 892 datasets) of the *A. thaliana* strains analyzed (Figures 1G, S2E), suggesting that differential methylation, and likely silencing, of *HISN6B* is not unique to Col-0. However, the basis for the differential methylation of *HISN6B* among strains is unclear.

In Col-0, homozygous *hisn6a-2* progeny of *hisn6a-2/HISN6A* heterozygotes arrest as pre-globular embryos (31). To test whether *HISN6B* derepression in the *hda6-7* mutant background rescues *hisn6a-2* lethality, we crossed a *hisn6a-2* heterozygote (-/+) with homozygous *hda6-7* (Figure 2A). Half of the resulting F1 plants harbor one *hisn6a-2* mutant allele (red ‘a’), one functional *HISN6A* allele (green ‘A’), one silent *HISN6B* allele from the *hisn6a-2* parent (red ‘B’), and one active, derepressed *HISN6B* allele (green ‘B’) from the *hda6-7* parent (Figure 2A). Among 89 of their F2 progeny, 17 (19%) *hisn6a-2* homozygous mutant (-/-) plants were recovered (Figure 2B, red bars). By contrast, zero *hisn6a-2* (-/-) plants were among the 64 progeny of self-fertilized *hisn6a-2*(-/+) plants wild-type for HDA6 (Figure 2B, black bars). Whereas wild-type Col-0 plants only express *HISN6A*, and *hda6-7* mutants express both *HISN6A* and *HISN6B*, the viable *hisn6a-2* homozygotes express only *HISN6B* (Figure 2C, *hisn6a-2;HISN6B*^active^).

**Figure 2.**
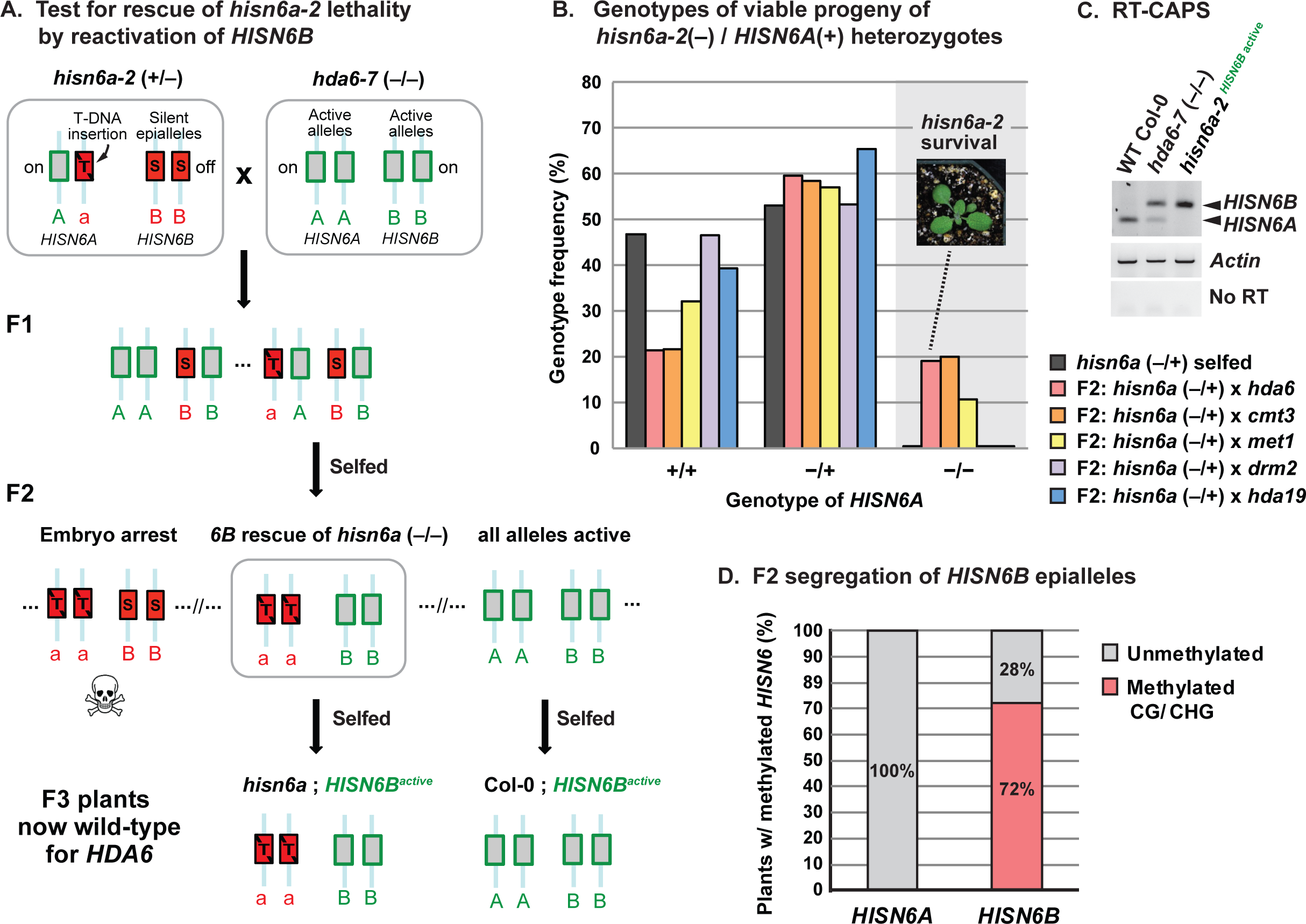
Reactivating *HISN6B* via elimination of symmetric DNA methylation rescues *hisn6a-2* lethality. **(A)** Strategy for synthetic rescue of *hisn6a-2* lethality. Red lowercase ‘a’ indicates the *hisn6a-2* null mutant allele, red uppercase ‘B’ indicates transcriptionally silent *HISN6B*, and green uppercase ‘A’ or ‘B’ indicates transcriptionally active *HISN6A* or *HISN6B*, respectively. *HDA6* genotypes are omitted for simplicity **(B)** Tests of *hisn6a-2* rescue by passage of *HISN6B* alleles through null mutants affecting gene silencing. Heterozygous *hisn6a-2* (+/-) was selfed (black), or crossed to *hda6-7* (pink), *cmt3-11t* (orange), *met1-7* (yellow), *drm2-2* (purple) or *hda19-1t* (blue). Selfed *hisn6a-2* (+/-) progeny (n=64 plants), or F2 progeny resulting from the crosses to other mutants (n=89, 60, 28, 60 and 30 plants, respectively) were genotyped for *hisn6a-2* and *HISN6A* alleles, whose frequencies were plotted. **(C)** RT-CAPS analysis of *HISN6A/B* gene expression in the *hda6-7* mutant and in a synthetically rescued *hisn6a-2* mutant line now harboring active *HISN6B* alleles. Actin reactions serve as loading controls. Reactions omitting reverse transcriptase (no RT) serve as controls for genomic DNA contamination. **(D)** Mendelian segregation of hypomethylated *HISN6B* epialleles. F2 progeny of *hisn6a-2* × *hda6-7* were assayed for the presence (red) or absence (grey) of cytosine methylation at the *Hpa*II/*Msp*I restriction site of *HISN6A/B* promoters. The percentage of plants in each category is plotted (n = 32). Unlike *HISN6B*, *HISN6A* did not show significant levels of promoter methylation (see Figures S3A and C).

We also crossed *hisn6a-2* (-/+) to the cytosine methyltransferase mutants, *cmt3-11t* or *met1-7*. Viable *hisn6a-2* homozygotes represented 20% of the F2 plants from the *cmt3-11t* cross and 11% of the F2 plants resulting from the *met1-7* cross (Figure 2B). By contrast, crosses to *drm2-2* or *hda19-t1*, a histone deacetylase that is functionally distinct from HDA6 (41, 42), yielded no viable *hisn6a-2* homozygous F2 progeny (Figure 2B). We conclude that *HISN6B* can be converted from a silent epiallele to an active allele in *hda6*, *cmt3* or *met1* mutants, allowing the re-animated gene to rescue plants lacking the normally essential *HISN6A* gene.

Analysis of F2 individuals in the *hisn6a-2* (-/+) × *hda6-7* segregating population revealed *HISN6B* promoter methylation in 72% of the plants, consistent with a 3:1 ratio due to Mendelian inheritance of methylated or unmethylated alleles (Figure 2D, S3A). Because the unmethylated *HISN6B* allele segregated independently of the *hda6-7* mutation, we were able to identify an F3 line (Col-0^*HISN6B* active^) that is homozygous for active *HISN6B* epialleles in an otherwise genetically wild-type background (Figures 2A; F3 individual #82 in Figure S3B). After self-fertilization of this line for three generations, no resetting to the silent, methylated state was observed (Figure S3C). This line, designated Col-0^*HISN6B* active^ was thus used for further genetic comparisons to wild type plants, designated Col-0^*HISN6B* silent^, homozygous for silent, methylated *HISN6B* epialleles. Analysis of methylome data obtained by Schmitz et al. (43) shows that methylated *HISN6B* epialleles were stably inherited over a span of 30 generations (Figure S3D).

In Cvi-0 × Col-0 F2 hybrids, homozygosity for both Col-0 *HISN6B* and Cvi-0 *HISN6A* is lethal (Figure 3A)(30). We confirmed this among 229 Cvi-0^*hisn6a* deletion^ × Col-0^*HISN6B* silent^ F2 progeny, using PCR to detect Col-0 *HISN6A* (abbreviated as A^C^), Col-0 *HISN6B* (B^C^), Cvi-0 *hisn6a* (a^i^) and Cvi-0 *HISN6B* (B^i^) alleles. Although ~14 individuals (^1^/16) could be expected to have the genotype a^i^a^i^ B^C^B^C^, none were observed (Figure 3B, arrow), a significant deviation (p = 4.8 × 10^-8^, Pearson’s chi-square test) indicating hybrid incompatibility. RT-CAPS analyses revealed that hybrid individuals that are homozygous for wild-type Col-0 (B^C^B^C^) *HISN6B* alleles fail to express *HISN6B*,whereas individuals inheriting at least one *HISN6B* allele from Cvi-0 (B^i^ or B^i^B^i^) showed *HISN6B* expression (Figure 3C). This indicates that silent or active *HISN6B* alleles of the parental strains are faithfully transmitted. The promoter methylation marks of Col-0 *HISN6B* (B^C^) alleles were also faithfully inherited among F2 hybrid individuals (Figure S4A).

**Figure 3.**
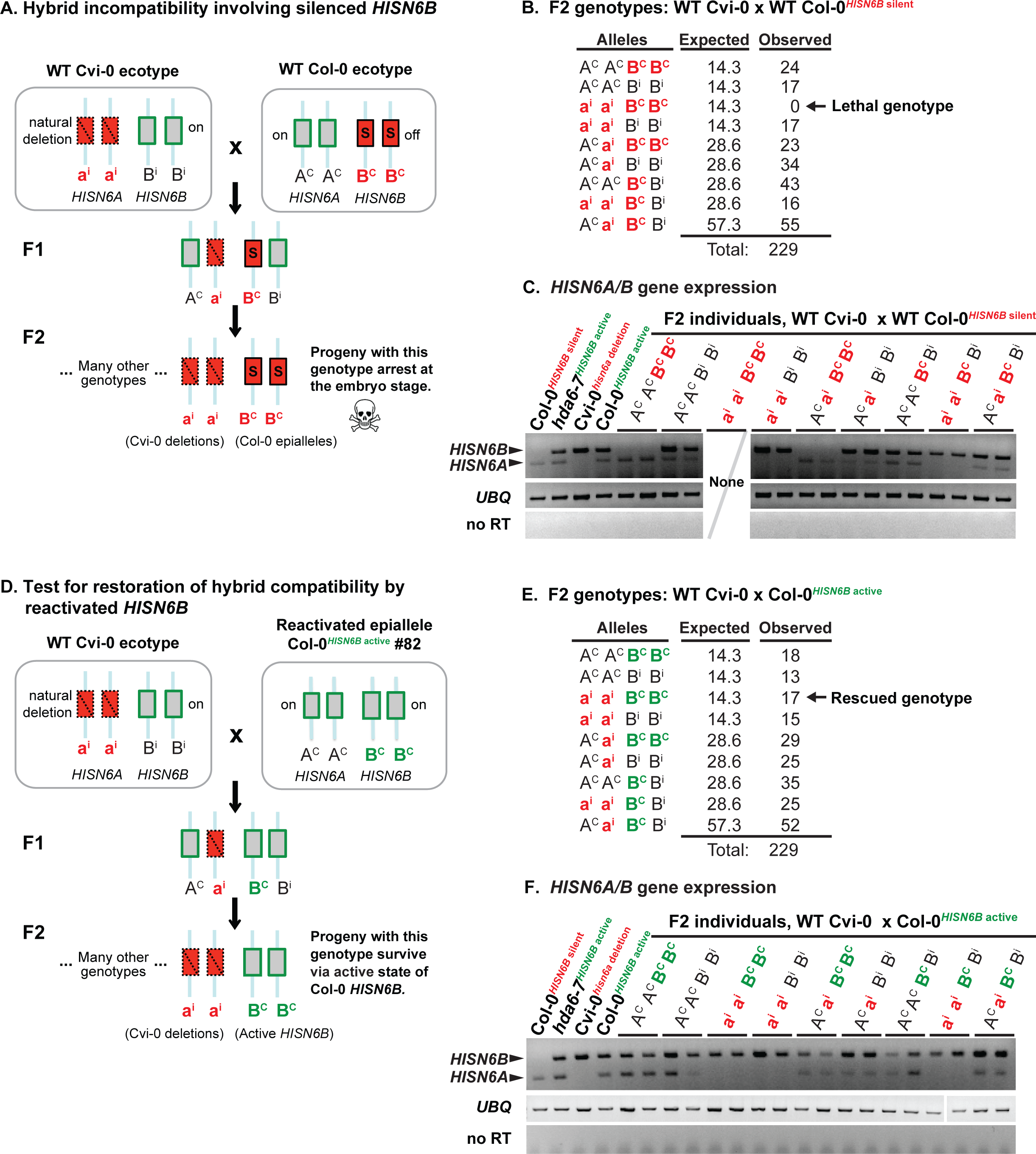
Active *HISN6B* epialleles circumvent lethality in *A.thaliana* Cvi-0 × Col-0 hybrids. **(A)** Schematic diagram of hybrid incompatibility between *A. thaliana* strains Col-0 and Cvi-0. Red ‘a^i^’ indicates Cvi-0 *hisn6a* deletion alleles, red ‘B^C^’ indicates silent Col-0 *HISN6B*, and black ‘A^C^’ or ‘B^i^’ indicate active Col-0 *HISN6A* or Cvi-0 *HISN6B*, respectively. Note that alleles inherited from Col-0 carry the superscript C, alleles from Cvi-0 carry the superscript i. All combinations of *HISN6A* and *HISN6B* alleles result in viable plants, except the double homozygous combination of Cvi-0 *HISN6A* alleles (a^i^a^i^) and Col-0 *HISN6B* alleles (B^C^B^C^). **(B)** F2 genotypes resulting from the cross: wild-type (WT) Cvi-0 × WT Col-0 (with silent *HISN6B*). A total of 229 individuals were genotyped. Expected allele frequencies were calculated assuming Mendelian segregation. Comparison of observed to expected frequencies using Pearson’s chi-square test resulted in a p-value of 4.8 × 10^-8^, indicating hybrid incompatibility between a^i^ a^i^ and B^C^B^C^ alleles. **(C)** RT-PCR analysis of *HISN6A/B* gene expression in F2 progeny of the WT Cvi-0^*hisn6a* deletion^ × WT Col-0^*HISN6B* silent^ cross. Two individuals for each of the 8 observed genotypes were assayed. Ubiquitin (UBQ) amplification products serve as loading controls. Reactions without reverse transcriptase (no RT) serve as controls for DNA contamination. Col-0^*HISN6B* silent^, Cvi-0^*hisn6a* deletion^, *hda6-7* and Col-0 carrying reactivated *HISN6B* alleles serve as *HISN6* expression controls. **(D)** Reactivated Col-0 *HISN6B* rescues Cvi-0 compatibility. The schematic depicts the genetic mechanism underlying rescue of hybrid compatibility following reanimation of Col-0 *HISN6B* alleles. Red ‘a^i^’ indicates Cvi-0 *hisn6a* deletion alleles, red ‘B^C^’ indicates silent Col-0 *HISN6B*, green ‘B^C^’ indicates active Col-0 *HISN6B*, and black ‘A^C^’ or ‘B^i^’ indicate active Col-0 *HISN6A* or Cvi-0 *HISN6B*, respectively. Note that alleles inherited from Col-0 carry the superscript C, alleles from Cvi-0 carry the superscript i. All combinations of *HISN6A* and *HISN6B* alleles result in viable plants, including the previously lethal combination of two Cvi-0 *HISN6A* alleles (a^i^a^i^) and two Col-0 *HISN6B* alleles (B^C^B^C^). **(E)** F2 genotypes of progeny resulting from the cross of WT Cvi-0 × Col-0 (active *HISN6B*). Asterisk indicated Pearson’s Chi-square (p-value = 0.78). Allele frequencies among 229 F2. Expected allele frequencies were calculated assuming Mendelian segregation. Comparison of observed and expected stats using Pearson’s chi-square test resulted in a p-value of 0.78, indicating hybrid compatibility has been restored. **(F)** *HISN6A/B* expression in the F2 progeny of the Cvi-0^*hisn6a* deletion^ × Col-0^*HISN6B* active^ cross. Two individuals from each of the 9 observed genotypes were assayed. UBQ reactions serve as loading controls. Reactions without reverse transcriptase (no RT) serve as controls for DNA contamination. Col-0^*HISN6B* silent^, Cvi-0^*hisn6a* deletion^, *hda6-7* and Col-0 carrying reactivated *HISN6B* serve as controls.

We next tested whether derepression of Col-0 *HISN6B* alleles would now allow the survival of F2 hybrids homozygous for both Col-0 *HISN6B* and Cvi-0 *HISN6A*. Indeed, the Cvi-0^*hisn6a* deletion^ × Col-0^*HISN6B* active^ cross yielded 17 healthy a^i^a^i^ B^C^B^C^ individuals in an F2 population of 229 plants (Figure 3E), which is statistically indistinguishable from the expected ~14 plants (p = 0.78, Pearson’s chi-square test). Col-0 *HISN6B* allele expression in the F2s (Figure 3F) correlates with the near absence of *HISN6B* methylation (Figure S4B). Collectively, these genetic tests show that reversion of *HISN6B* epialleles from a methylated, silent state to a hypomethylated, active state eliminates *HISN6*-based hybrid incompatibility between the Col-0 and Cvi-0 strains of *A. thaliana*.

## Discussion

Our study shows that strain-specific silencing of duplicated *HISN6* genes occurs in *A. thaliana* and is maintained by symmetric CG and CHG methylation involving HDA6, MET1 and CMT3. MET1 and CMT3-dependent DNA methylation can maintain silent epialleles over numerous meiotic generations, independent of initial silencing signals (15, 16, 21-23, 44). Mutation of *MET1*, *CMT3* or *HDA6* converts silent *HISN6B* epialleles into active, unmethylated alleles that are stably transmitted according to Mendelian rules of segregation. Moreover, reactivation of *HISN6B* circumvents the normal lethality of *hisn6a* mutations in Col-0 and prevents the occurrence of *HISN6*-based hybrid lethality among Cvi-0 × Col-0 hybrid progeny.

Hybrid incompatibility involving *HISN6A* and *HISN6B* alleles fits the model of Lynch and Force in that alternative *HISN6A* versus *HISN6B* expression states lead to deleterious gene combinations in Cvi-0 × Col-0 F2 progeny (7, 30). Unlike the Bateson, Dobzhansky and Muller model, Lynch and Force’s model for hybrid incompatibility does not require gene neo-functionalization. Instead, differential loss of function of one member of a duplicated gene pair in different subpopulations or strains spawns incompatibilities if the strains hybridize. Examples in plants include the *DPL1* and *DPL2* genes of *O. sativa* subspecies *indica* and *japonica* (45), and *AtFOLT1* and *AtFOLT2* genes of *A. thaliana* (28). The latter report showed that Col-0 × C24 and Col-0 × Sha incompatibilities correlate with the duplication, and additional complex rearrangement, of *AtFOLT1* in the C24 and Sha strains. These mutations trigger methylation of *AtFOLT1*, leaving *AtFOLT2* as the only active copy in C24 or Sha, and causing hybrid incompatibility with Col-0, which lacks *AtFOLT2* altogether. Our study demonstrates that transgenerationally heritable (but fully revertible) epialleles that have not undergone tandem duplication, rearrangement or mutation also contribute to hybrid incompatibility, providing additional evidence that epigenetic variation can foster reproductive isolation (28, 46, 47) in a manner consistent with the Lynch and Force hypothesis.

## Materials & Methods

### Plant materials

*Arabidopsis thaliana* strain Col-0 used in the study was a Pikaard lab stock. Strain Cvi-0 (CS22614) was obtained from the Arabidopsis Biological Resource Center at Ohio State University. *hda6-6* (a.k.a., *axe1-5*) and *hda6-7* (a.k.a., *rts1-1*) mutants were described in Murfett et al. (2001)(37) and Aufsatz et al. (2002)(36). *hda19-1t* is the *athd1-t1* (Ws) mutant allele of Tian et al. (2003)(41) introgressed into Col-0 by three backcrosses. The RNA polymerase mutants *pol IV* (*nrpd1-3*) and *pol V* (*nrpe1-11*) were described in Onodera et al. (2005)(48) and Pontes et al. (2006)(49). The mutants *drm2-2, cmt2-3* (SALK_012874), *cmt3-11t* (SALK_148381), *hisn6a-2* (SAIL_750) and *hisn6a-3*(SALK_089516), and triple mutant *drm1-2 drm2-2 cmt3-11t* were obtained from the Arabidopsis Biological Resource Center. *met1-3* was described in Saze et al. (2003)(15) and *met1-7* (SALK_076522) was obtained from the Nottingham Arabidopsis Stock Center.

### RNA analyses

Total RNA was extracted from two-week-old rosettes or from inflorescences using TRIreagent (MRC, Inc.). For semi-quan
titative RT-PCR, 1.5 μg of DNase I-treated total RNA was used for random-primed cDNA synthesis by Superscript III reverse transcriptase (Invitrogen). Standard PCR was performed on cDNA aliquots (~100 ng RNA input) using GoTaq Green (Promega) and primers listed in **Table S1**. PCR products were analyzed either directly by agarose gel electrophoresis (*Actin* and *UBQ* controls) or following restriction enzyme digest (*RsaI*) and then agarose gel electrophoresis (RT-CAPS for *HISN6A/B*).

### DNA analyses

For genotyping, genomic DNA was purified from two-week-old seedlings using a CTAB extraction protocol. GoTaq Green master mix (2X, Promega) was mixed with ~100 ng of genomic DNA and particular genotyping primer pairs. PCR products were scored either directly by agarose gel electrophoresis or following restriction enzyme digest (*Bsp*HI) and then agarose gel electrophoresis (CAPS). For DNA methylation analyses, genomic DNA was isolated from inflorescence tissue using the Nucleon PhytoPure DNA extraction kit (Amersham). Chop PCR assays were performed using 100 ng of restriction enzyme-digested (“chopped”) genomic DNA as in Earley et al. (2010)(50). Primers used for genotyping and Chop PCR are listed in **Table S1**.

### Bioinformatic analyses

Analysis of wild-type and mutant methylomes of strain Col-0 (Figure 1E, S2C) were performed on data from NCBI GEO GSE39901 (6 datasets, wiggle format)(39). Methylation profiles were based on *HISN6A* gene model AT5G10330.8 and *HISN6B* gene model AT1G71920.2, whose similar intron/exon structures facilitated comparison of the two genes. Fractional cytosine methylation in CG, CHG and CHH sequence contexts were converted from wiggle to bigWig format, extracted using the bwtool software and then plotted in Microsoft Excel(51). For the comparison of *A.thaliana* ecotype methylomes (Figures 1G, S2E), datasets were obtained from NCBI GEO GSE43857 (927 ecotype datasets, tabular format)(40). After removing redundant or unidentified ecotypes, 892 methylomes remained for analysis. For each methylome, CG, CHG and CHH methylation were separately tallied over 100-bp, non-overlapping windows in *HISN6A* or *HISN6B*, respectively, starting 500 bp upstream of the transcription start site (+1) and stopping 300 bp downstream of the transcription termination site (TTS). Hierarchical clustering was then performed using Euclidean distance and Ward’s method to regroup ecotypes with similar patterns of cytosine methylation along *HISN6A* or *HISN6B*. Heatmaps were drawn using the “heatmap.2” function of R. For *HISN6B*, ecotypes were assigned to two visually evident categories: methylated promoter versus unmethylated promoter.

### Statistical tests

Chi-square goodness-of-fit tests were performed on genotype data in Figures 3B and 3E to test the hypothesis of *HISN6B* methylation-dependent hybrid incompatibility and generate p-values in each case. The GPower software(52) estimates that sample sizes of at least 158 F2 plants would be needed to determine whether the observed allele frequencies deviate from Mendelian expectations, which are far exceeded by the 229 plants genotyped in each experiment.

## Acknowledgements

TB and JW designed the study and performed all experimental procedures. TB and DP performed bioinformatic analyses of *HISN6A/B* gene methylation using publically available methylome data. FP assisted in RNA-seq analyses of *hda6* mutants. TB and CSP wrote the manuscript. CSP is an Investigator of the Howard Hughes Medical Institute (HHMI) and Gordon and Betty Moore Foundation (GBMF). This work was supported by National Institutes of Health grant GM077590 and GBMF grant GBMF3036 to CSP. TB was supported by an NIH Ruth L. Kirschstein National Research Service Award, and funding from HHMI.

## Supplementary Information

### Supplemental Figure and Table Legends

**Table S1.**
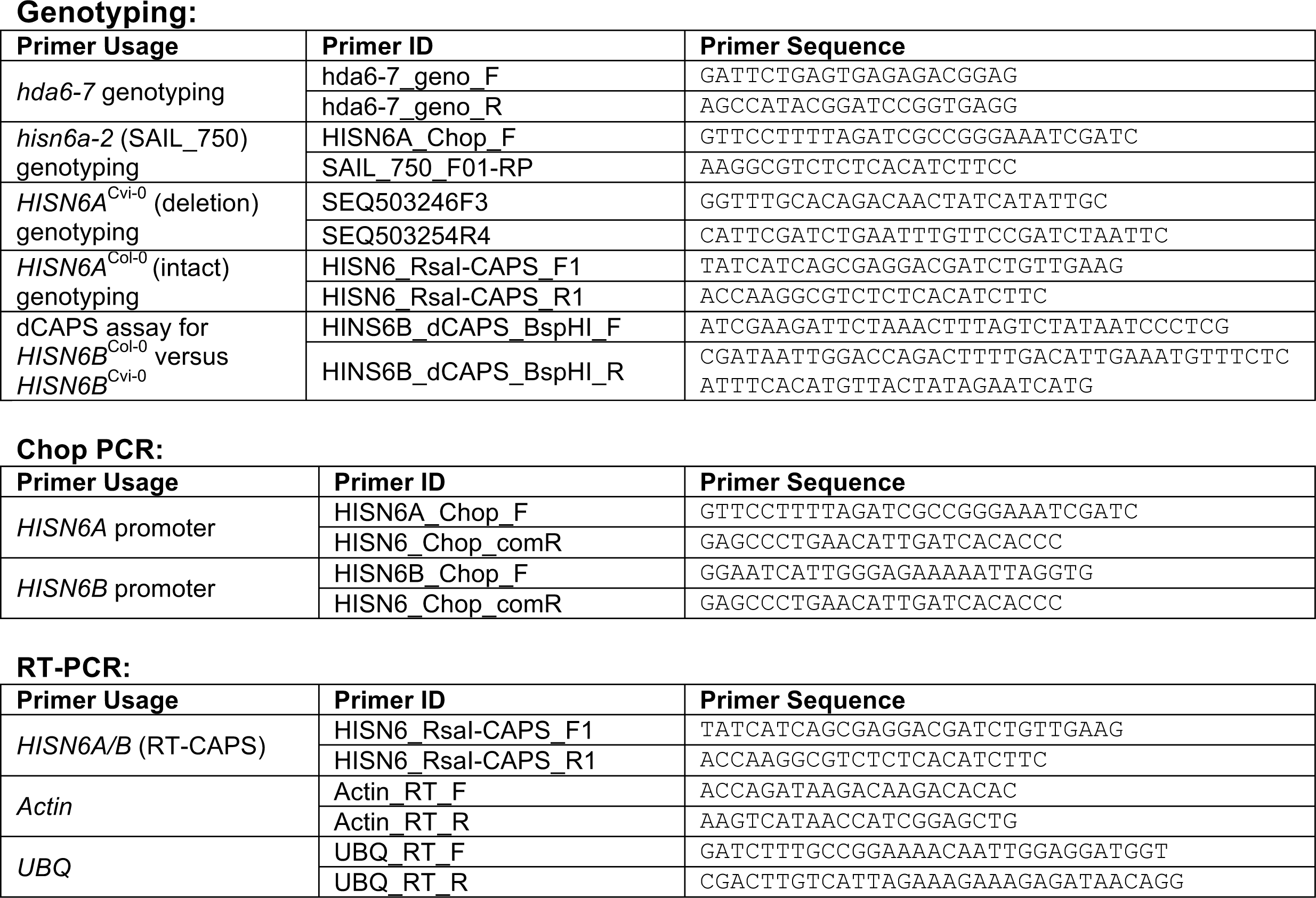
Oligonucleotides used for genotyping, Chop-PCR and RT-PCR

**Figure S1.**
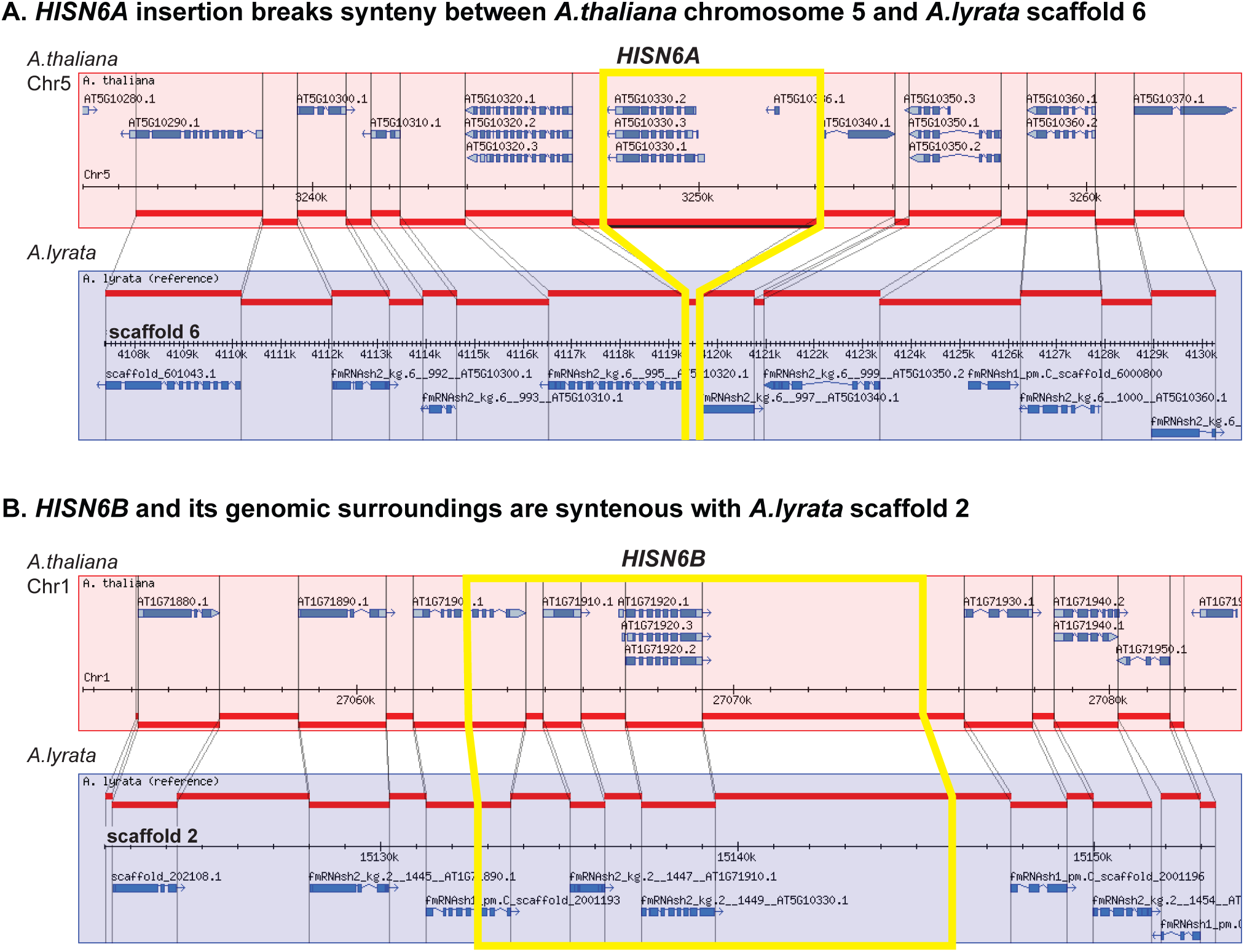
Chromosomal contexts of *HISN6A* and *HISN6B*. **(A)** The *HISN6A* locus of *A. thaliana* interrupts a region displaying extensive synteny between *A. thaliana* Chr 5 and *A. lyrata* scaffold 6, suggesting insertion at this site after the divergence of the two species from a common progenitor (see also, Ingle et al., (32). **(B)** *HISN6B* is present within a region displaying extensive synteny with *A. lyrata* scaffold 2, the location of the sole *HISN6* gene of *A. lyrata*.

**Figure S2.**
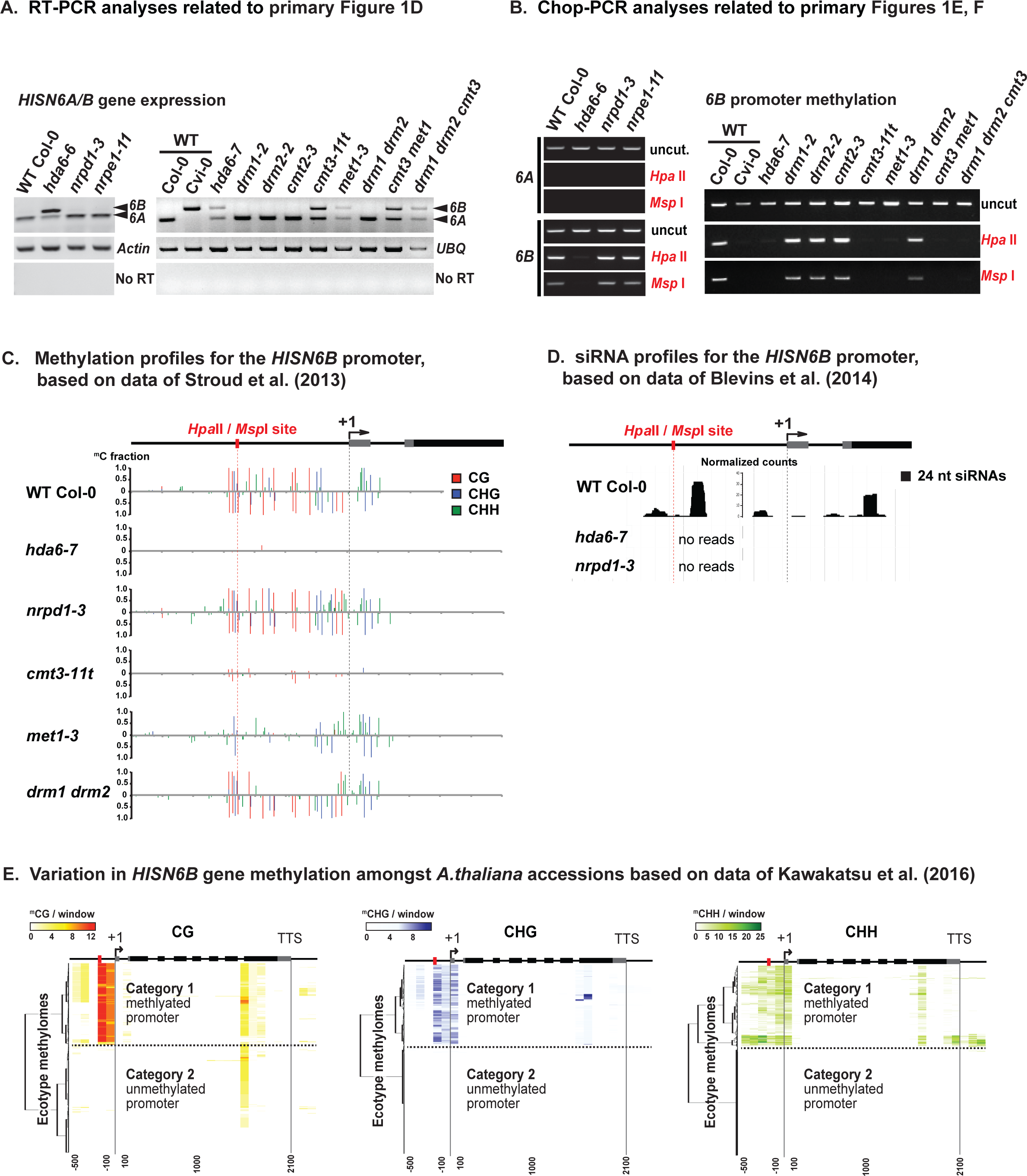
*HISN6A* and *HISN6B* expression and methylation in wild-type Col-0 and in mutants affecting chromatin modification and gene silencing. **(A)** RT-PCR products were digested with *RsaI* to discriminate *HISN6A/B* expression in WT Col-0 or Cvi-0 and the indicated Col-0 mutants, as in Figure 1D. (**B)** *HISN6A* and *HISN6B* promoter methylation in wild-type and various Col-0 mutants was analyzed by Chop-PCR at the promoter region *HpaII* / *MspI* restriction site, as in Figures 1E and 1F. Reactions omitting restriction enzymes (uncut) control for equivalent DNA input in all reactions. **(C)** Cytosine methylation profiles are shown for a 600-bp region encompassing the *HISN6B* promoter and transcription start site (+1). Methylomes of 3-week-old leaves isolated from WT and mutant lines of the Col-0 strain were analyzed (data from Stroud et al., 2013). Bars represent fractional methylation per cytosine position, color-coded by sequence context (CG, CHG and CHH). The *HpaII* / *MspI* restriction enzyme site used for Chop PCR assays is highlighted (red dotted line). **(D)** A genome browser shot showing small RNA profiles of 2-week-old leaves of WT, *hda6-7* or *nrpd1-3* mutants of the Col-0 strain. Read counts corresponding to 24-nt siRNAs are plotted in black on the y-axis, normalized by total mapped reads (data from Blevins et al., 2014) (14). **(E)** Hierarchical clustering analysis of *HISN6A/B* methylation, as in Primary Figure 1G, but divided by CG, CHG or CHH sequence context (data from Kawakatsu et al., 2016).

**Figure S3.**
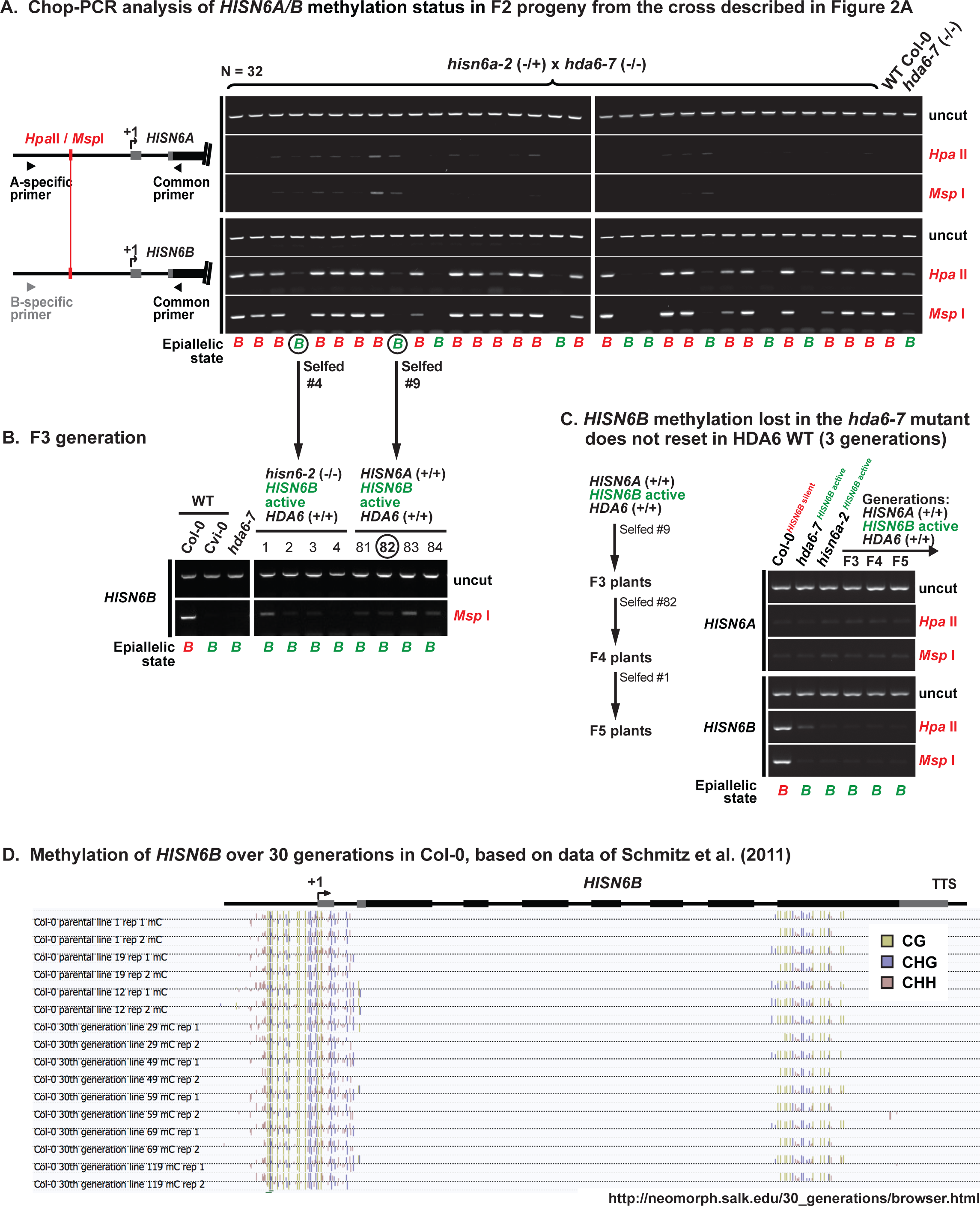
Stable *HISN6B* methylation is lost in *hda6* mutants and not regained upon restoration of wild-type *HDA6* genes. **(A)** Chop-PCR DNA methylation analysis of F2 progeny from the cross in Figure 2A testing *HISN6A/B* methylation at the promoter *Hpa*II/ *Msp*I site. Reactions omitting restriction enzymes serve to control for equivalent DNA input to all reactions (uncut). PCR specificity was verified by sequencing uncut PCR products. **(B)** Chop-PCR analysis of of *HISN6A/B* promoter methylation in F3 progeny from the cross in Figure 2A. **(C)** Chop-PCR analysis of *HISN6A/B* promoter methylation in a *HISN6B* active lineage derived from an *hda6* mutant background, in three self-fertilized generations after restoration of HDA6 function (F3, F4 and F5 generation from the cross in Figure 2A). WT Col-0 (*HISN6B* silent) and *hda6-7* and *hisn6a-2* mutants (both *HISN6B* active) serve as controls. **(D)** Browser shot of *HISN6B* methylation profiles in WT Col-0 lines propagated by single-seed descent for 30 generations. Profiles of independent parental lines and 30th generation lines are shown, with methylcytosine sequence context (CG, CHG or CHH) indicated using different colors. The data are from the study of Schmitz et al. (2011), displayed using the genome browser tool described in that paper: http://neomorph.salk.edu/30_generations/browser.html).

**Figure S4.**
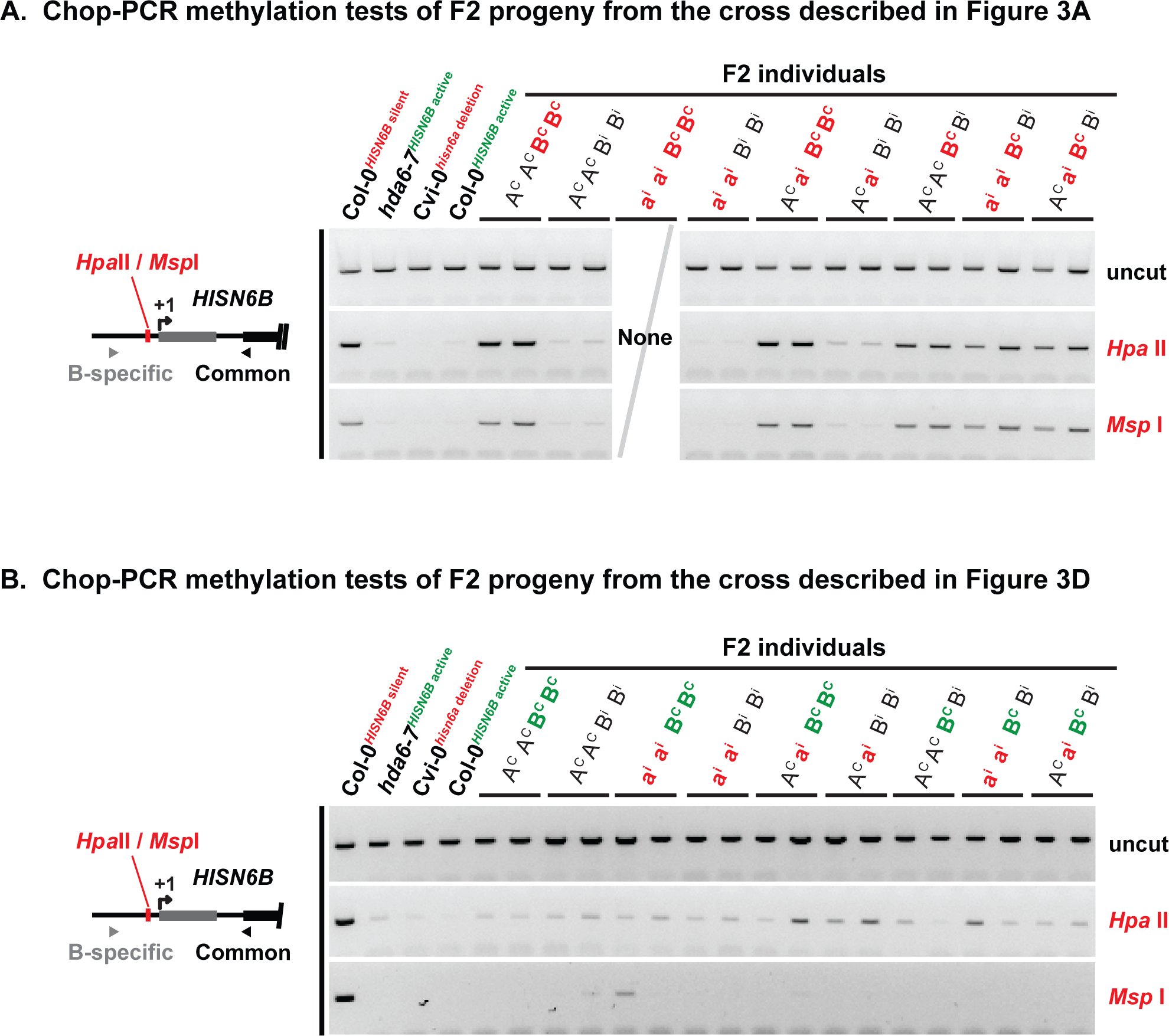
Methylation status of *HISN6B* alleles inherited from wild-type Col-0 or Cvi-0, or a Col-0 *hda6-7* mutant. **(A)** Chop-PCR DNA methylation analysis of F2 progeny from the cross in Figure 3A, assessing methylation at the promoter region *HpaII*/*MspI* site. The doubly homozygous combination: Cvi-0 *HISN6A* (a^i^a^i^) and Col-0 *HISN6B* (B^C^B^C^) was not recovered in N=229 plants and was thus not available for analysis. Reactions omitting restriction enzymes (uncut) reveal equivalent DNA input in all reactions. Primer specificity was verified by sequencing uncut PCR products. **(B)** Chop-PCR analysis of F2 progeny from the cross in Figure 3D.

